# Vibrational behavior of psyllids (Hemiptera: Psylloidea): functional morphology and mechanisms

**DOI:** 10.1101/593533

**Authors:** Yi-Chang Liao, Zong-Ze Wu, Man-Miao Yang

## Abstract

Vibrational behavior of psyllids was first documented more than six decades ago. Over the years, workers have postulated as to what the exact signal-producing mechanisms of psyllids might be but the exact mechanism has remained elusive. The aim of this study is to determine the specific signal-producing structures and mechanisms of the psyllids. Here we examine six hypotheses of signal-producing mechanisms from both previous and current studies that include: wing vibration, wing-wing friction, wing-thorax friction, wing-leg friction, leg-abdomen friction, and axillary sclerite-thorax friction. Through selective removal of possible signal producing structures and observing wing-beat frequency with a high-speed video recorder, six hypotheses were tested. Extensive experiments were implemented on the species *Macrohomotoma gladiata* Kuwayama, while other species belonging to different families, i.e., *Trioza sozanica* (Boselli), *Mesohomotoma camphorae* Kuwayama, *Cacopsylla oluanpiensis* (Yang), and *Cacopsylla tobirae* (Miyatake) were also examined to determine the potential prevalence of each signal-producing mechanism within the Psylloidea. Further, scanning electron microscopy (SEM) was used to examine possible rubbing structures. The result of high speed photography showed that wing-beating frequency did not match the dominant frequency of vibrational signals, resulting in the rejection of wing vibration hypothesis. As for the selective removal experiments, the axillary sclerite-thorax friction hypothesis is accepted and wing-thorax friction hypothesis is supported partially, while others are rejected. The SEM showed that the secondary axillary sclerite of forewing bears many protuberances that would be suitable for stridulation. In conclusion, the signal-producing mechanism of psyllids involves two sets of morphological structures. The first is stridulation between the axillary cord and anal area of the forewing. The second is stridulation between the axillary sclerite of the forewing and the mesothorax.

## Introduction

Acoustic communication is prevalent among the insects with more than 18 orders having been recorded to communicate via acoustic signals [1]. Acoustic signals of insects usually play an important role in mating behavior [2–4]. Other functions of acoustic signals also include defense [5] and food searching [6–8]. In Hemiptera, the mechanisms of sound producing vary greatly, for example, members of Heteroptera (bugs) can emit signals by stridulation, tymbal buckling, or abdomen vibration, while many species of Auchenorrhyncha (cicadas, planthoppers, leafhoppers, treehoppers) utilize a tymbal organ to produce signals [9]. Additionally, whiteflies (Sternorrhyncha, Aleyrodoidea) produce signals through abdominal oscillation [10] and aphids (Aphidoidea) emit sound by rubbing the abdomen and hind legs against substrates [11]. As well, psyllids (Psylloidea) are comparatively active singers during their mating behavior but until now their mechanisms have been poorly understood [12–16].

The vibrational behavior of psyllids was first reported by Ossiannilsson in 1950 [17]. In this work, he suggested that psyllids produced faint signals via wing vibration. Two years later, Tuthill [18] suggested that the radula of the forewing may hold the potential for stridulation. Later that same year, Heslop-Harrison [19] proposed a novel sounding mechanism of psyllids in which stridulation occurred between the inner side of legs and the bee-hive-like structure of the abdomen. He also suggested that there was potentially a second sound producing mechanism of psyllids, via leg-leg stridulation and through examination of members of 85% of the known psyllid genera from all geographic areas, found that psyllids did not possess a tymbal-like structure [20]. Some two decades later, Taylor [21] suggested psyllids make signals by stridulation between the anterior margin of forewing and axillary cord on the mesoscutellum and metascutellum. As well, Tishechkin [22] examined the thorax structure of 14 species among four families and agreed with the hypothesis of Taylor [21]. Wenninger et al. [23] stated that psyllids make signals only via wing vibration and without any stridulation. More recently, Eben et al. [24] agreed that psyllids produce signals by stridulation between the axillary cord of the thorax and anal area of the forewings. Despite all of the work and hypotheses of others in the past and partly due to the small size and difficult observation of these taxa, the exact mechanism has remained speculative and unclear.

Based on previous studies of the potential sound producing mechanisms of psyllids, we have identified six hypotheses to be examined. The description of these six hypotheses are as follows: (1) wing vibration hypothesis [17, 23]: psyllids make signals via wing vibration without any friction between structures; (2) wing-wing friction hypothesis [18, 20]: psyllid forewings possess tooth-like structure so as to rub each other when psyllids vibrate their wings; (3) wing-leg friction hypothesis [20]: this hypothesis was also proposed by Heslop-Harrison in the same reference; (4) wing-thorax friction hypothesis [12, 21, 22, 24]: forewing rubs against the thorax when wings vibrate. Specifically, friction occurs between the anal area of forewing and axillary cord located on mesoscutellum and metascutellum; (5) leg-abdomen friction hypothesis [19]: abdomen contraction can be seen when psyllids emit signals, therefore, Heslop-Harrison thought that the friction occurs between inner side of leg and abdomen, of which, the first sternite possess a rough bee-hive-like structure; (6) axillary sclerite-thorax friction hypothesis: this is a new hypothesis that is first proposed in this study. We suggest here that the axillary sclerite makes contact with the thorax while the forewings vibrate rapidly and that is how psyllids produce vibrational signals.

As mentioned above, most hypotheses are involved with wings, hence, specific experiments are designed to verify each hypothesis and examine the most likely mechanism of psyllid sound production. These experiments are implemented through wing-cut and observation, through using high-speed video recorder to record the numbers of wing vibrations when psyllids emit signals and by taking scanning electron microscopy (SEM) images to observe the existence of specific anatomy that might serve as sound producing structures.

## Materials and methods

### Insect preparation

Psyllid larvae were collected from the field (Table 1) and positioned in an incubator maintained in a dark environment and temperature at 25±2°C. Adults do not mate at night so the lack of light was utilized to maintain virginity [25]. Larvae were raised within a plastic box (27 × 19 × 5 cm) in which plant shoot cuttings were inserted in a vial of water. Emerging adults were isolated into a small plastic box (15 × 8 × 5 cm) and sexed immediately after eclosion.

**Table 1.**
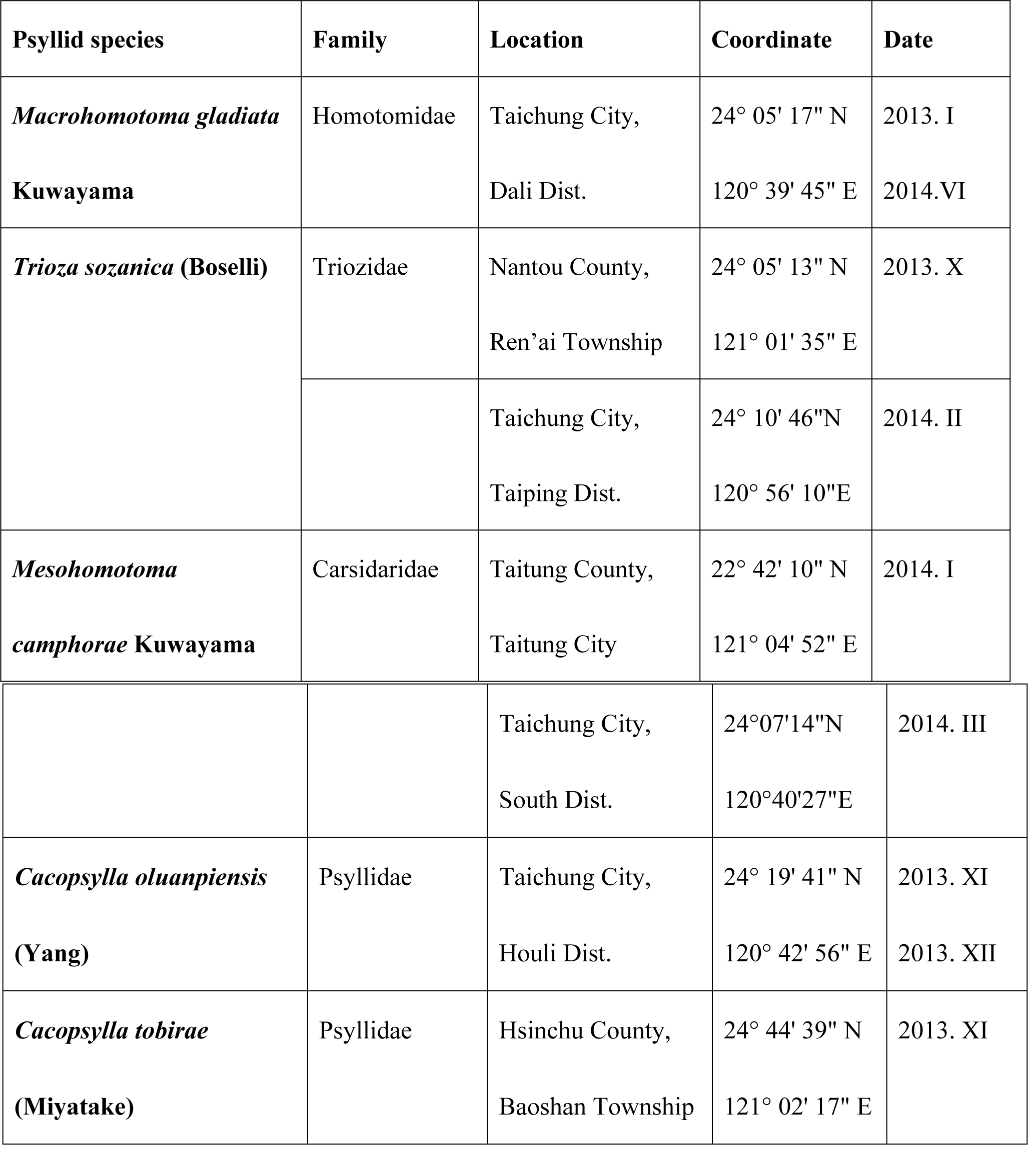
Collecting information of psyllid specimens for the experiments on potential sound producing mechanisms.

Extensive experiments were implemented on the species *Ma*. *gladiata* Kuwayama. This species was originally widespread in the Orient and South Asia but has become an invasive pest in Europe and USA more recently [26–30]. The vibrational behavior of this species has been described in detail [15]. Body size of *Ma*. *gladiata* is comparatively large and this species actively makes duet signals, which makes it a good model species to be observed and conduct serial experiments on.

In the mating behavior of *Ma*. *gladiata*, the male produces a two-chirp calling and the female replies with a single chirp, after that, the male makes a single chirp in response to the female then searches for her [15]. As vibrational signals of psyllids differ among different taxa even in closely related species, different species may possess different sound producing behavior [13–15]. Other species belonging to different families, i.e., *T*. *sozanica* (Boselli), *Me*. *camphorae* Kuwayama, *C*. *oluanpiensis* (Yang), and *C*. *tobirae* (Miyatake) were also examined to determine the prevalence of signal-producing and the associated mechanisms within the Psylloidea. *Me*. *camphorae* (Carsidaridae) is widespread in low elevations of Taiwan and feeds on *Hibiscus* and *Urena* spp. (Malvaceae); *T*. *sozanica* (Triozidae) is a pit gall inducer that is specific to *Daphniphyllum* spp. (Daphniphyllaceae); *C*. *oluanpiensis* (Psyllidae) and *C*. *tobirae* feed on *Pittosporum tobira* Ait. and *Pittosporum pentandrum* (Blanco) Merr., respectively. The latter two species of *Cacopsylla* have been described in detail for their species properties and vibrational behavior [14].

### Recording of vibrational signals

Vibrational recording of psyllids mainly follows the methods of Liao and colleagues [13, 14]. Recording was conducted in an anechoic chamber illuminated with a 2-foot long flurorescent tube. In each trial, a single psyllid individual (usually male) was gently settled on a plant shoot which was inserted into a water vial. The plant shoot was confined in a plastic tube (15 × 5 cm) to restrict the psyllids from jumping away. Signals emitted from the psyllids were received by a gramophone stylus which slightly touched the base of the plant shoot and was then amplified through an amplifier (Lzban, DRA-455, China) and saved in a dictaphone (Laxon, USB-F 20, Taiwan). If psyllids did not make any signals for a while, we used a dictaphone (Sony NWZ-E435, Japan) to playback recorded conspecific signals to induce its vibrational behavior. Sampling rate for signal recording was 48,000 Hz.

### Wing-cut experiments

We designed a series of wing-cut experiments to examine the role of wings in producing signals (Fig 1). The treatments are described below. A1: without any treatment (control); A2: forewing cut; A3: hindwing cut; A4: both wings cut; A5: anal area of forewing cut; A6: forewing cut but axillary sclerite left. Adults were immobilized by chilling them on a cold pack. Specific parts of wings were removed by a customized scissor. Psyllids that had been treated were settled in the same rearing area for at least one night before recording was attempted.

**Fig 1.**
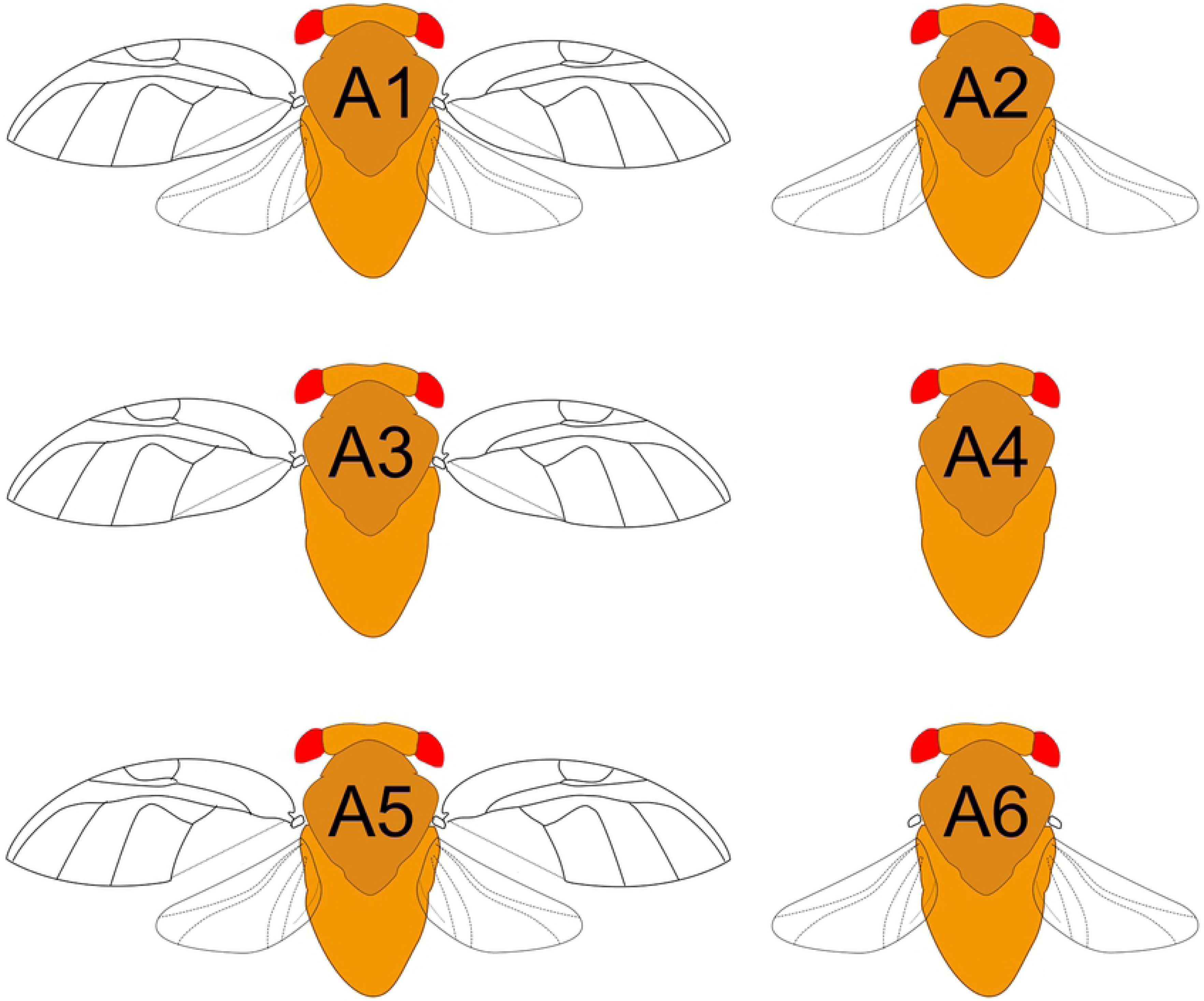
Illustration of each treatment of wing-cut in the study. A1: Control (entire forewing and hindwing); A2: Forewing cut; A3: Hindwing cut; A4: Both wings cut; A5: Anal area of forewing cut; A6: Forewing cut but axillary sclerite left.

*Macrohomotoma gladiata* was used in all six treatments. After verification of the sound producing mechanism of *Ma*. *gladiata*, other psyllid species belonging to different families were used to confirm the prevalence of certain mechanisms. We selected six individuals from the other four species representing 3 families for further examination, i.e., *T*. *sozanica, Me*. *camphorae*., *C*. *oluanpiensis*, and *C*. *tobirae*. First, we recorded their original signals as a control treatment (A1). After that, three individuals of each species were treated by cutting the forewing (A2), while the other three individuals were treated by cutting the forewing but leaving the axillary sclerite intact (A6). Then, we recorded the vibrational signals of the individuals of A2 and A6 of these four species again and compared their signals with A1.

### Wing vibration recording

A high-speed video recorder (GC-PX100BU, JCV, Canada) equipped with 8X microlens was positioned and focused on an adult to observe their call. The tested individuals were the same as control, namely A1 treatment. This experiment was only completed on *Ma*. *gladiata*. Through high-speed video recorder (600 frames per second), we counted the number of *Ma. gladiata* wing vibrations and from that were able to determine the frequency.

### Statistical analysis and plotting

When possible, we picked up two to three vibrational signals from a single individual for analysis. The signal processing and statistical analysis was conducted by using Matlab 8.0 (R2012b, Mathworks, Natick, MA). For file reading we used scripts by Ellis [31]. The script for noise reduction and plotting was modified from Vincent [32] and Zhivomirov [33]. We picked up the maximum voltage ratio of every signal after noise reduction to represent the amplitude of signals. The signal amplitude of psyllids from different treatments was compared using Wilcoxon signed-rank test and Kruskal-Wallis test by Matlab 8.0 with Statistics Toolbox 8.1. Plot was made via Sigma Plot 10.0 (Systat Software, San Jose, CA).

### Scanning electron microscopy

Psyllid specimens previously used for recordings of vibrational signals were preserved in 70% ethanol. The specimens were kept in 42 degree oven for at least 10 hours to ensure they were dried out. Each individual was dissected to examine the dorsal and lateral view of the thorax and the right axillary sclerite. Also to facilitate this, the head, wings, legs, and abdomen were also removed. Specimens were mounted on standard card stock and coated with gold via a sputter coater (Polaron SC502, UK). All images were taken with a scanning electron microscope (SEM) (Topcon ABT-150S, Japan) located at the Department of Plant Pathology, National Chung Hsing University, Taichung, Taiwan.

## Results

### The wing-beat frequency

With the aid of the high-speed video recorder, we measured the wing-beat frequency of males of *Ma*. *gladiata*. The first chirp was 118.32 ± 5.68 Hz and the second chirp was 139.97 ± 4.18 Hz. The second chirp always having a higher wing vibration per second.

### Wing-cut treatments

According to the control treatment (A1) of *Ma*. *gladiata*, the dominant frequency of first chirp was 805.4 ± 186.0 Hz and the second chirp was 932.0 ± 187.6 Hz. Among the six treatments, the individuals of A1, A3, A5, and A6 were able to make sound, however, A2 and A4 could not (Table 2). The amplitude of voltage ratio of A1 (0.730 ± 0.183) and A3 (0.673 ± 0.151) were significantly higher than that of A5 (0.395 ± 0.226) and A6 (0.245 ± 0.132) (Fig 2).

**Table 2.**
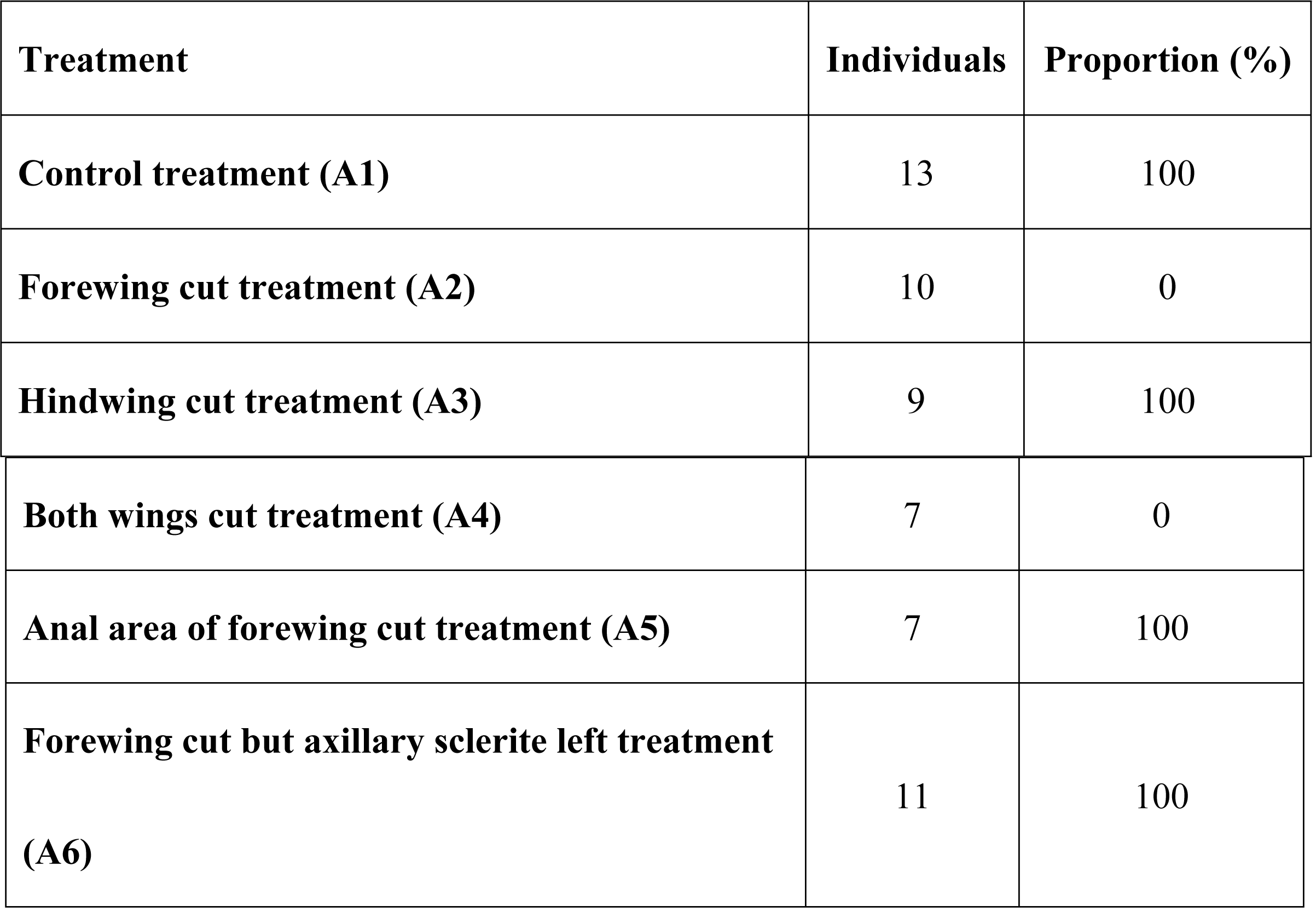
Relationship between wing area and sound production of *Macrohomotoma gladiata* Kuwayama.

| Treatment | Individuals | Proportion (%) |
| --- | --- | --- |
| Control treatment (A1) | 13 | 100 |
| Forewing cut treatment (A2) | 10 | 0 |
| Hindwing cut treatment (A3) | 9 | 100 |
| **Both wings cut treatment (A4)** | 7 | 0 |
| **Anal area of forewing cut treatment (A5)** | 7 | 100 |
| **Forewing cut but axillary sclerite left treatment (A6)** | 11 | 100 |

**Fig 2.**
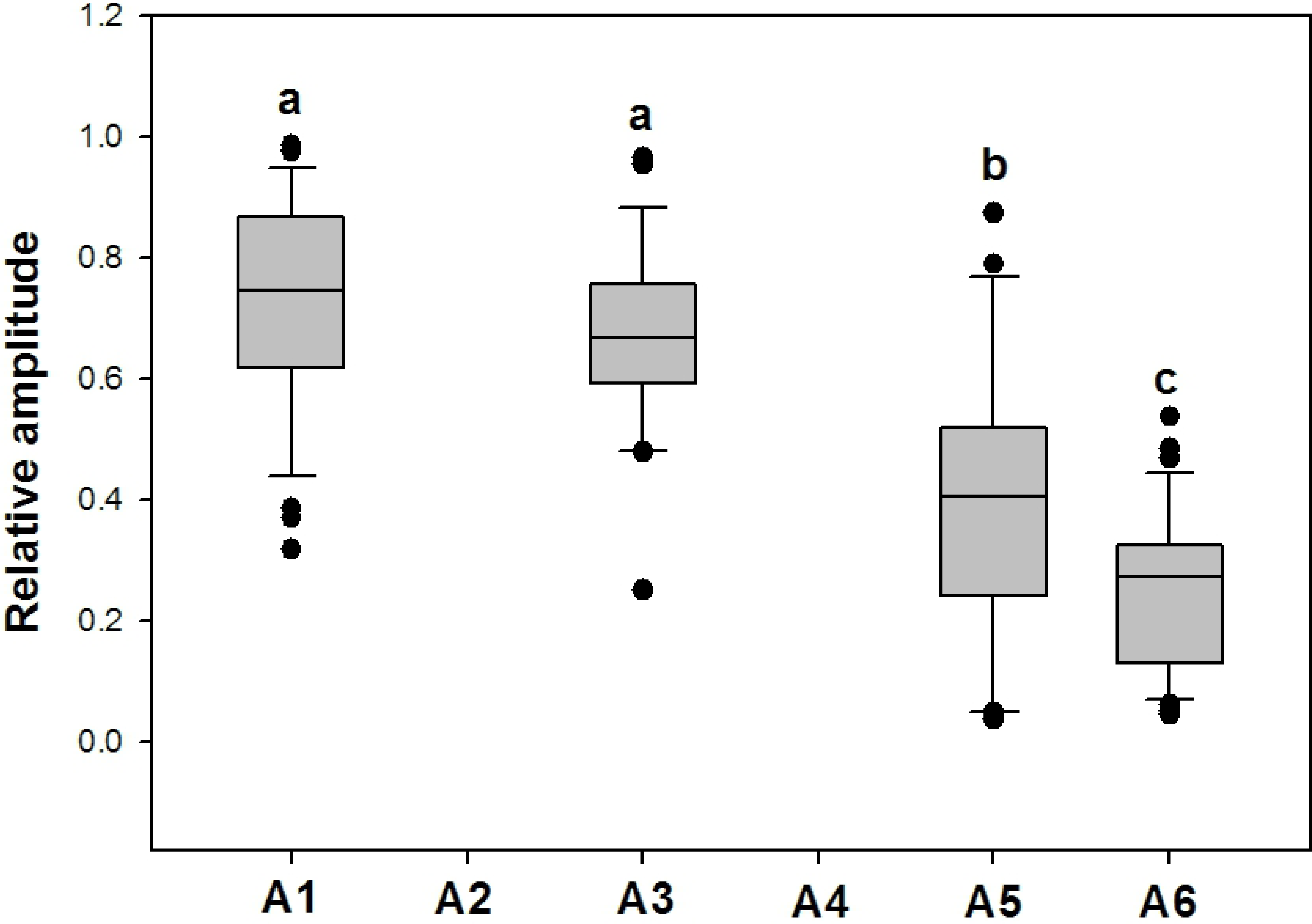
Signal amplitude of male calling of *Macrohomotoma gladiata* in each treatment. A1: Control (entire forewing and hindwing); A2: Forewing cut; A3: Hindwing cut; A4: Both wings cut; A5: Anal area of forewing cut; A6: Forewing cut but axillary sclerite left. (Kruskal Wallis test, H = 70.73, *P* < 0.001)

The results for the other four species of psyllids, i.e., *Me*. *camphorae, T*. *sozanica, C*. *oluanpiensis*, and *C*. *tobirae*, were consistent with that of *Ma*. *gladiata*. Individuals of both A1 and A6 could make sound, however, A2 could not make sound (Table 3). The amplitude of volume ratio of A1 were significantly larger than that of A6 in these four psyllid species (Table 3).

**Table 3.**
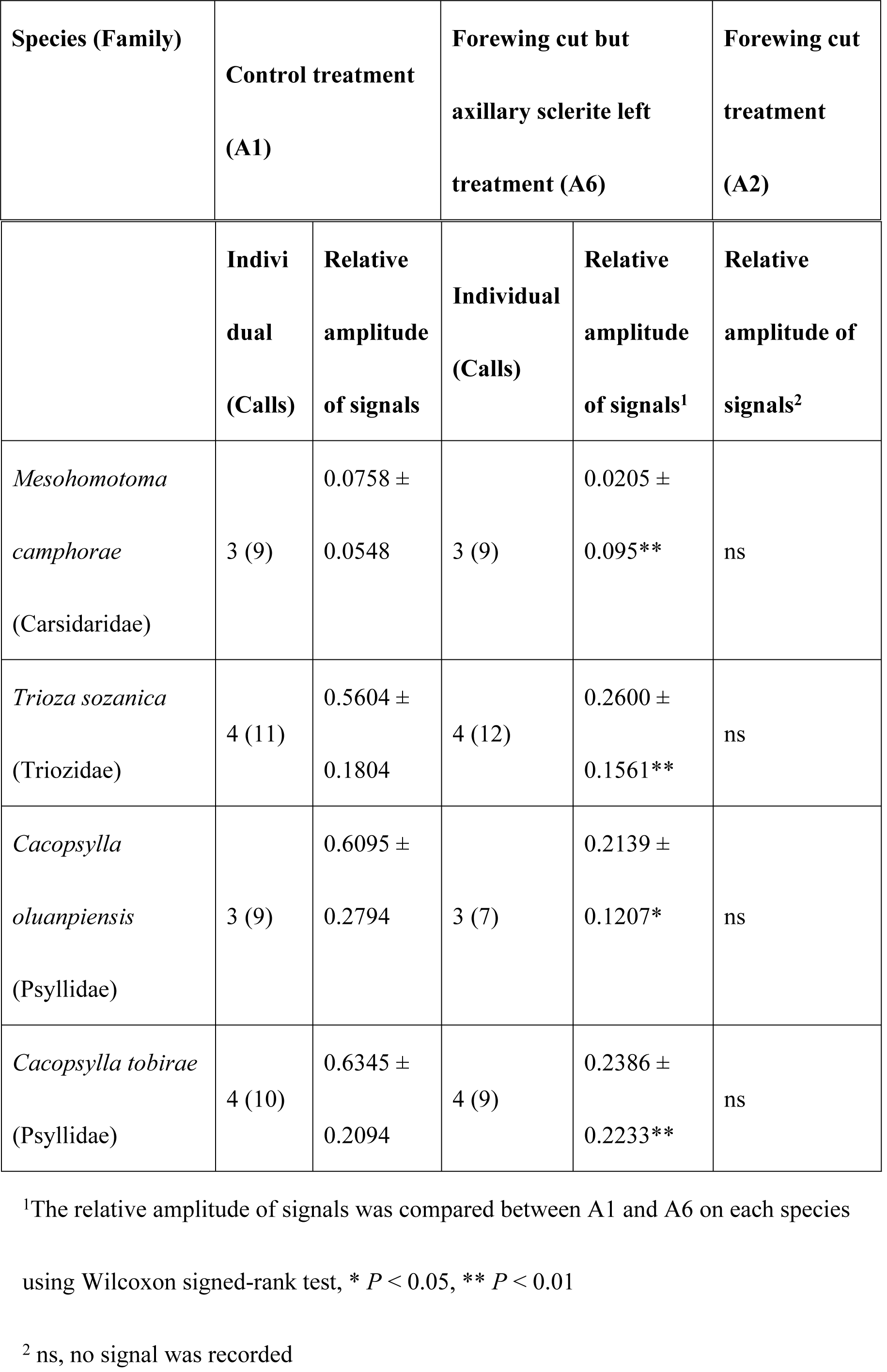
The effect of forewing cut (A2) and forewing cut but axillary sclerite left treatment (A6) on signal amplitude in psyllid species belonging to different families.

### The surface of the psyllid sound making structure

With the cutting treatments and sound records in mind, it seems probably that the axillary sclerite is the main area for psyllids to produce signals. SEM images of the dorsal and lateral view of *Ma. gladiata, Me*. *camphorae, T*. *sozanica, C*. *oluanpiensis*, and *C*. *tobirae* show that the dorsal surface of secondary axillary sclerite is rough and possesses many protuberances (Fig 3). In addition, the rough surface on mesothorax appears to form the area of friction with the dorsal surface of the secondary axillary sclerite (Fig 3).

**Fig 3.**
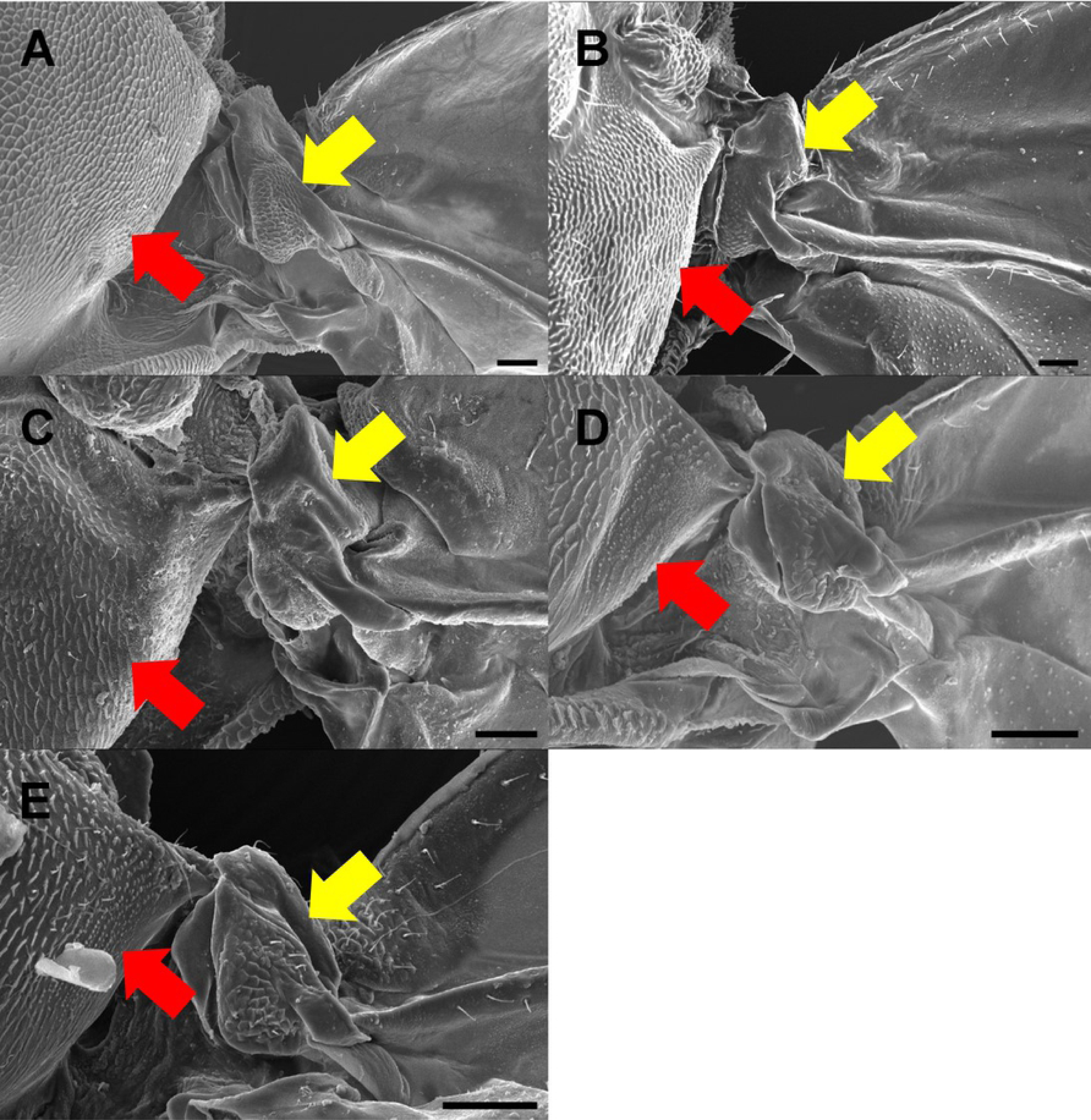
Photographs of scanning electron microscopy showing dorsal view of axillary sclerite in five speices of psyllids belonging to different families. **A:** *Macrohomotoma gladiata* Kuwayama; B: *Trioza sozanica* (Boselli); C: *Mesohomotoma camphorae* Kuwayama; D: *Cacopsylla oluanpiensis* Yang; E: *Cacopsylla tobirae* Miyatake. Yellow arrow: Protuberances on the surface of secondary axillary sclerite. Red arrow: scale-like surface of mesothorax scutum.

## Discussion

As a result of the wing-cut experiments, we were able to elucidate the crucial relationship between sound making and wings in the treatments of A1, A2, A3, and A4. Consequently, according to forewing cut (A2) and hindwing cut treatments (A3), we ensured that psyllids are not able to make sound without forewing and it is now clear that the hindwing is certainly not evolved in sound production. The psyllids with only forewings left were able to produce sound, and the signal amplitude of those individuals was not significantly different compared with control treatment (Fig 2; Table 3). According to this evidence, we confirm the importance of the forewing in contributing to the sound producing mechanism of psyllids.

### Hypothesis testing

#### Wing vibration hypothesis

Wing vibration of insects such as mosquitos and bees, can produce rarefaction waves, which are transmitted through the air [34]. Usually the dominant frequency of these signals is consistent with the frequency of the wing-beat [35, 36]. In this study, the dominant frequency of vibrational signals and frequency of wing beat did not match between dominant frequency and wing beat and there is about seven-fold difference. It is also not likely that the dominant frequency we recorded is a seventh harmonic of fundamental frequency of wing-beat because the amplitude of the signal drops drastically at higher harmonics [37]. Moreover, the major transmitted route for psyllid signals is the substrate, i.e. plant twigs and leaves, not the air. Thus, we confirmed that psyllids could not make sound only by wing vibration, which means that friction must occur. This hypothesis is rejected.

#### Wing-wing friction hypothesis

Individuals from the A2 treatments were not able to produce signals. This suggests that the hindwing is not involved in sound producing mechanism of psyllids. As well, individuals from the A6 treatments are able to produce signals which also supports rejection of this hypothesis. In addition, through the observation with high-speed video recording, we can see that the forewing movement is vertical when psyllids produce signals. Wing-wing friction is likely to be horizontal in nature as seen in the sound producing of male crickets.

#### Wing-thorax friction hypothesis

This hypothesis suggested that friction between axillary cord of the thorax and anal area of the forewing is responsible for sound production. Based on morphological examination in previous studies, all psyllids have an axillary cord on their thorax [21, 22]. However, according to the results from forewing cut (A2) and both wings cut treatments (A4), psyllids could not make sound without their forewings. This also further supports our belief that the hindwing has no influence on sound producing. Moreover, based on the hindwing cut treatment (A3), we have confirmed the importance of the forewing for sound production. Results of forewing anal area cut treatment (A5) showed the mean amplitude of signal was half compared with that of the control treatment. This suggests that anal area of forewing plays a considerably important role on sound production. Although this result does not fit the hypothesis completely, it does show the significance of the anal area of the forewing for sound production.

#### Leg-abdomen friction hypothesis

During psyllid calling, we can usually see the upward movement and slight vibration of the abdomen of psyllids. For example, the abdomen of *T*. *sozanica*, which produces a long call duration (> 10 seconds), would move upward at the beginning of producing signals and move downward when calling ended. Further, we did not observe obvious rubbing behavior between the leg and abdomen when psyllids produce signals, suggesting that this hypothesis is likely to be invalid.

#### Wing-leg friction hypothesis

Heslop-Harrison [20] thought the wing-leg hypothesis was the minor mechanism of sound producing and that leg-abdomen friction was the major mechanism of sound production in psyllids. However, the leg-abdomen friction hypothesis was rejected by this study based on the hindwing cut treatment (A3) and forewing cut in combination with the left axillary sclerite treatment (A6), because individuals of these two treatments were able to produce sound.

#### Axillary sclerite-thorax friction hypothesis

This hypothesis was first raised in this study. Individuals from the anal area of forewing cut treatment (A5) were able to make sound signals. This suggests that there may be other structures related to sound producing involved. The axillary sclerite of the forewing is a heavily-scleritized structure that appears to have the potential for producing signals and is the reason for proposing this hypothesis. We noticed that individuals of the forewing cut (A2) and both wings cut treatments (A4) were not able to produce signals; however, individuals with the forewing cut but the axillary sclerite left intact (A6) were able to produce signals suggesting that this theory may have some validity. The amplitude of signals of the forewing cut with axillary sclerite intact (A6) are significantly lower than that of hindwing cut (A3) and control treatments (A1).

Based on examination of SEM photographs, the sclerotized structure of the axillary sclerite in psyllids were rough and may be suitable for friction with the thorax to produce signals. Especially the second axillary sclerite which is well-developed structurally and possesses scale-like structure on the surface.

### The sound producing mechanism of psyllids

We conclude that the sound producing mechanism of Psylloidea has two components. One is via the stridulation between anal area of forewing and the axillary cord on mesothorax. The other is via stridulation between axillary sclerite of forewing and thorax. The finding of the axillary sclerite as a sound producing mechanism is a novel scientific finding. The forewings of psyllids play an important role during sound producing but the loss of the hindwing does affect sound production (Fig 2). Results showed that the signal amplitude declines if the anal area of forewing was removed or only the axillary sclerite is left intact. This phenomenon suggest that the remigium of the forewing may serve as an amplifier of signals.

### Sound producing structure and systematics

Taxonomic usefulness of acoustic apparati in insects, is well documented. Villet [38] used several characters of the tymbal organ in his taxonomic revisions, including shape and size of operculum, as well as the meracanthus, and ribs on the tymbal. In Orthoptera, the teeth number on the file of the forewing of crickets are different among species and they serve as diagnostic characters [39]. It is now apparent that the morphology of the acoustic apparatus of psyllids may be used in species identification as well. Tishechkin [22] pointed out that the axillary cord is different among psyllids and suggested that axillary cord possesses characteristics distinguishable at the family level. This study has uncovered that the scale-like structure on the axillary sclerite of psyllids can be further measured and compared and appears to be useful taxonomically.

The current accepted phyogeny of Hemiptera suggests that the Sternorrhyncha is an older clade within Hemiptera [40]. Three superfamilies of Sternorrhyncha, Psylloidea, Aphidoidea, and Aleyrodoidea, have various sound producing mechanisms. This indicated that the sound producing mechanism of each group may evolved independently as they do not possess a tymbal-like structure for sound production. Members of Auchenorrhyncha and Heteroptera, such as Aphrophoridae, Cicadellidae, Dictyopharidae, Issidae, Alydidae, Cydnidae, Rhopalidae, produce signals using a tymbal [9]. This phenomenon suggests that the tymbal mechanism is a synapomorphic character that evolved after Sternorrhyncha branched off. The sound producing mechanism of Psylloidea confirmed by this study may be an apomorphy for this group. This finding provides evidence of the potential monophyly of Psylloidea. Also, we found that axillary sclerites were different in scale pattern among species under SEM and this structure may have the potential to be taxonomically useful at the species level. Furthermore, quantitative analysis for characteristics of the axillary sclerite could be further conducted for potential use in delineating the higher classification of Psylloidea.

## Acknowledgments

We would like to thank the following colleagues in National Chung Hsing University: Chun-I Chiu and Yin-Hsuan Teng for providing advise on the wing-cut experiments; Dr. Kai-Jung Chi for her assistance in application of high-speed video recording; Yu-Chun Lin for her help in modification of an early draft; Dr. Jeng-Tze Yang for providing the anechoic chamber; and Wesley Hunting for his kind help in English editing.

## References

1. Cocroft RB, Rodríguez RL. The behavioral ecology of insect vibrational communication. BioScience. 2005; 55: 323–334. doi: 10.1641/0006-3568(2005)055[0323:TBEOIV]2.0.CO;2

2. Bell PD. Multimodal communication by the black-horned tree cricket, Oecanthus nigricornis (Walker) (Orthoptera: Gryllidae). Can J Zool. 1980; 58: 1861–1868. doi: 10.1139/z80-254

3. Hoy RR, Hoikkala A, Kaneshiro K. Hawaiian courtship songs: evolutionary innovation in communication signals of Drosophila. Science. 1988; 240:217–219. doi: 10.1126/science.3127882

4. Doherty J. Song recognition and localization in the phonotaxis behavior of the field cricket, Gryllus bimaculatus (Orthoptera: Gryllidae). J Comp Physiol A. 1991; 168: 213–222. doi: 10.1007/BF00218413

5. Holman J. Possible sound producing structures present in some Macrosiphini (Homoptera: Aphididae). Eur J Entomol. 1994; 91:97–101.

6. Cocroft RB. Vibrational communication facilitates cooperative foraging in a phloem-feeding insect. Proc R Soc B Biol Sci. 2005; 272: 1023–1029. doi: 10.1098/rspb.2004.3041

7. Evans TA, Lai JCS, Toledano E, McDowall L, Rakotonarivo S, Lenz M. Termites assess wood size by using vibration signals. PNAS. 2005; 102: 3732–3737. doi: 10.1073/pnas.0408649102

8. Ulyshen MD, Mankin RW, Chen Y, Duan JJ, Poland TM, Bauer LS. Role of Emerald Ash Borer (Coleoptera: Buprestidae) larval vibrations in host-quality assessment by Tetrastichus planipennisi (Hymenoptera: Eulophidae). J Econ Entomol. 2011; 104: 81–86. doi: 10.1603/EC10283

9. Virant-Doberlet M, Cokl A. Vibrational communication in insects. Neotrop Entomol. 2004; 33: 121–134. doi: 10.1590/S1519-566X2004000200001

10. Kanmiya K, Sonobe R. Records of two citrus pest whiteflies in Japan with special reference to their mating sounds (Homoptera: Aleyrodidae). Appl Entomol Zool. 2002; 37: 487–495. doi: 10.1303/aez.2002.487

11. Kubota S. Rubbing behaviours in some aphids. Jpn J Entomol. 1985; 53: 595–603.

12. Percy DM, Taylor GS, Kennedy M. Psyllid communication: acoustic diversity, mate recognition and phylogenetic signal. Invertebr Syst. 2006; 20: 431–445. doi: 10.1071/is05057

13. Liao YC, Huang SS, Yang MM. Substrate-borne signals, specific recognition, and plant effects on the acoustics of two allied species of Trioza, with the description of a new species (Psylloidea: Triozidae). Ann Entomol Soc Am. 2016; 109: 906–917. doi: 10.1093/aesa/saw060

14. Liao YC, Yang MM. Acoustic communication of three closely related psyllid species: a case study in clarifying allied species using substrate-borne signals (Hemiptera: Psyllidae: Cacopsylla). Ann Entomol Soc Am. 2015;108: 902–911. doi: 10.1093/aesa/sav071

15. Liao YC, Yang MM. First evidence of vibrational communication in Homotomidae (Psylloidea) and comparison of substrate-borne signals of two allied species of the genus Macrohomotoma Kuwayama. J Insect Behav. 2017; 30: 567–581. doi: 10.1007/s10905-017-9640-2

16. Yang MM, Yang CT, Chao J. Reproductive isolation and taxonomy of two Taiwanese Paurocephala species (Homoptera: Psylloidea). Taiwan Mus Spec Publ. 1986; 6: 176–203.

17. Ossiannilsson F. Sound production in psyllids (Hem. Hom.). Opus Entomol. 1950; 15: 202.

18. Tuthill LD. On the Psyllidae of New Zealand (Homoptera). Pac Sci. 1952; 6: 83–125.

19. Heslop-Harrison G. XXVII.—The number and distribution of the spiracles of the adult psyllid. Ann Mag Nat Hist. 1952; 5: 248–260.

20. Heslop-Harrison G. Sound production in the Homoptera with special reference to sound producing mechanisms in the Psyllidae. J Nat Hist Ser 13. 1960; 3: 633–640. doi: 10.1080/00222936008651067.

21. Taylor KL. A possible stridulatory organ in some Psylloidea (Homoptera). Aust J Entomol. 1985; 24: 77–80.

22. Tishechkin DY. On the structure of stridulatory organs in jumping plant lice (Homoptera: Psyllinea). Russian Entomol J. 2006; 15: 335–340.

23. Wenninger EJ, Hall DG, Mankin RW. Vibrational communication between the sexes in Diaphorina citri (Hemiptera: Psyllidae). Ann Entomol Soc Am. 2009; 102: 547–555.

24. Eben A, Mühlethaler R, Gross J, Hoch H. First evidence of acoustic communication in the pear psyllid Cacopsylla pyri L. (Hemiptera: Psyllidae). J Pest Sci. 2015; 88, 87–95. doi: 10.1007/s10340-014-0588-0

25. Wenninger EJ, Hall DG. Daily timing of mating and age at reproductive maturity in Diaphorina citri (Hemiptera: Psyllidae). Fla Entomol. 2007; 90: 715–722. doi: 10.1653/0015-4040(2007)90[715:DTOMAA]2.0.CO;2

26. Mifsud D, Porcelli F. The psyllid Macrohomotoma gladiata Kuwayama, 1908 (Hemiptera: Psylloidea: Homotomidae): a Ficus pest recently introduced in the EPPO region. EPPO Bulletin. 2012; 42: 161–164. doi: 10.1111/j.1365-2338.2012.02544.x

27. Bella S, Rapisarda C. New findings in Italy of the recently introduced alien psyllid Macrohomotoma gladiata and additional distributional records of Acizzia jamatonica and Cacopsylla fulguralis (Hemiptera Psylloidea). Redia. 2014; 97:151–155.

28. Laborda R, Galán-Blesa J, Sánchez-Domingo A, Xamaní P, Estruch VD, Selfa J, et al. Preliminary study on the biology, natural enemies and chemical control of the invasive Macrohomotoma gladiata (Kuwayama) on urban Ficus microcarpa L. trees in Valencia (SE Spain). Urban For Urban Gree. 2015; 14: 123–128. doi:http://dx.doi.org/10.1016/j.ufug.2014.12.007

29. Pedata PA, Burckhardt D, Mancini D. Severe infestations of the jumping plant-louse Macrohomotoma gladiata, a new species for Italy in urban Ficus plantations. B Insectol. 2012; 65: 95–98.

30. Rung A. A new pest of ficus in California: Macrohomotoma gladiata Kuwayama, 1908 (Hemiptera: Psylloidea: Homotomidae), new to North America. Check List. 2016; 12: 1–5. doi: 10.15560/12.3.1882

31. Ellis D. mp3read and mp3write [cited August 2010]. Available from: http://www.mathworks.com/matlabcentral/fileexchange/13852-mp3read-and-mp3write.

32. Vincent C. Vuvuzela sound denoising algorithm [cited August 2010]. Available from: http://www.mathworks.com/matlabcentral/fileexchange/27912-vuvuzela-sound-denoising-algorithm.

33. Zhivomirov H. Sound analysis with Matlab Implementation [cited August 2014]. Available from: http://www.mathworks.com/matlabcentral/fileexchange/38837-sound-analysis-with-matlab-implementation.

34. Unwin D, Corbet SA. Wingbeat frequency, temperature and body size in bees and flies. Physiol Entomol. 1984; 9: 115–121.

35. Williams CM, Galambos R. Oscilloscopic and stroboscopic analysis of the flight sounds of Drosophila. Biol Bull. 1950; 99: 300–307. doi: 10.2307/1538745

36. Webb JC, Sharp JL, Chambers DL, Benner JC. Acoustical properties of the flight activities of the caribbean fruit fly. J Exp Biol. 1976; 64: 761–772.

37. Arthur BJ, Emr KS, Wyttenbach RA, Hoy RR. Mosquito (Aedes aegypti) flight tones: frequency, harmonicity, spherical spreading, and phase relationships. J Acoust Soc Am. 2014; 135: 933–941. doi: 10.1121/1.4861233

38. Villet MH. The cicada genus Stagira Stål 1861 (Homoptera Tibicinidae): systematic revision. Trop Zool. 1997; 10: 347–392. doi: 10.1080/03946975.1997.10539347

39. Walker TJ, Carlysle TC. Stridulatory file teeth in crickets: Taxonomic and acoustic implications (Orthoptera: Gryllidae). Int J Insect Morphol Embryol. 1975; 4: 151–158. doi: 10.1016/0020-7322[75]90013-6

40. Cui Y, Xie Q, Hua J, Dang K, Zhou J, Liu X, et al. Phylogenomics of Hemiptera (Insecta: Paraneoptera) based on mitochondrial genomes. Syst Entomol. 2013; 38: 233–245. doi: 10.1111/j.1365-3113.2012.00660.x

